# Identification of a family of *Vibrio* type III secretion system effectors that contain a conserved serine/threonine kinase domain

**DOI:** 10.1101/2020.12.11.418251

**Authors:** N Plaza, IM Urrutia, K Garcia, MK Waldor, CJ Blondel

## Abstract

*Vibrio parahaemolyticus* is a marine Gram-negative bacterium that is a leading cause of seafood-borne gastroenteritis. Pandemic strains of *V. parahaemolyticus* rely on a specialized protein secretion machinery known as the type III secretion system 2 (T3SS2) to cause disease. The T3SS2 mediates the delivery of effector proteins into the cytosol of infected cells, where they subvert multiple cellular pathways. Here, we identify a new T3SS2 effector protein encoded by VPA1328 (VP_RS21530) in *V. parahaemolyticus* RIMD2210633. Bioinformatic analysis revealed that VPA1328 is part of a larger family of uncharacterized T3SS effector proteins with homology to the VopG effector protein in *V. cholerae* AM-19226. These VopG-like proteins are found in many but not all T3SS2 gene clusters and are distributed among diverse *Vibrio* species including *V. parahaemolyticus*, *V. cholerae*, *V. mimicus*, and *V. diabolicus* and also in *Shewanella baltica*. Structure-based prediction analyses uncovered the presence of a conserved C-terminal kinase domain in VopG orthologs, similar to the serine/threonine kinase domain found in the NleH family of T3SS effector proteins. However, in contrast to NleH effector proteins, in tissue culture-based infections, VopG did not impede host cell death or suppress IL-8 secretion, suggesting a yet undefined role for VopG during *V. parahaemolyticus* infection. Collectively, our work reveals that VopG effector proteins, a new family of likely serine/threonine kinases, is widely distributed in the T3SS2 effector armamentarium among marine bacteria.

**IMPORTANCE:** *Vibrio parahaemolyticus* is the leading bacterial cause of seafood-borne gastroenteritis worldwide. The pathogen relies on a type III secretion system to deliver a variety of effector proteins into the cytosol of infected cells to subvert cellular function. In this study, we identified a novel *Vibrio parahaemolyticus* effector protein that is similar to the VopG effector of *Vibrio cholerae.* VopG-like effectors were found in diverse *Vibrio* species and contain a conserved serine/threonine kinase domain that bears similarity to the kinase domain in the EHEC *and Shigella* NleH effectors that manipulate host cell survival pathways and host immune responses. Together our findings identify a new family of *Vibrio* effector proteins and highlight the role of horizontal gene transfer events among marine bacteria in shaping T3SS gene clusters.

## INTRODUCTION

*Vibrio parahaemolyticus* is a marine Gram-negative bacterium that is the leading bacterial cause of seafood-borne gastroenteritis worldwide (1). In 1996, a new clonal *V. parahaemolyticus* strain of the O3:K6 serotype, now known as the pandemic clone, emerged and has been responsible for major outbreaks of gastroenteritis in diverse locations around the globe (2).

In addition to the presence of the characterized virulence factors thermostable direct hemolysin (TDH) and the *tdh*-related hemolysin (TRH), genome sequencing revealed that all *V. parahaemolyticus* strains encode a type III secretion system on chromosome 1 (T3SS1) (3). Furthermore, strains related to the pandemic clone harbor an evolutionarily distinct T3SS known as T3SS2 (4–6) encoded within an 80kb *Vibrio parahaemolyticus* pathogenicity island 7 (VPaI-7) on chromosome 2 (3). T3SSs are multicomponent nanomachines that enable Gram-negative bacteria to deliver proteins known as effectors directly from the bacterial cytosol into the cytosol of eukaryotic cells. Translocation of effectors into host cells enables pathogens to hijack host-cell signaling, thereby manipulating a variety of host cell functions (reviewed in (7)). Indeed, the virulence of many human, animal, and plant pathogens depends on the activity of the T3SS injectisome and the repertoire of effector proteins delivered to their respective hosts’ cells (8, 9).

Notably, most *V. parahaemolyticus* strains isolated from human clinical samples harbor T3SS2 and studies in animal models have shown that T3SS2 is essential for *V. parahaemolyticus* to colonize the intestine and to cause enteritis and diarrhea (10–12). Therefore, T3SS2 is considered a key *V. parahaemolyticus* virulence factor. Several T3SS2 related gene clusters have been identified in other *Vibrio* species and are referred to as T3SS2 phylotypes (T3SS2α, T3SS2β and T3SS2γ) (13). T3SS2α include T3SS2 gene clusters related to those found in the *tdh*-positive *V. paraha*emolyticus pandemic strain RIMD2210633 and in *V. cholerae* strain AM-19226. T3SS2β include T3SS2 gene clusters related to those found in *V. parahaemolyticus* strain TH3996 and *V. cholerae* strain 1587 (14). Finally, T3SS2γ include T3SS2 gene clusters related to those encoded in *V. parahaemolyticus* strain MAVP-Q, which has features found in the T3SS2α and T3SS2β gene clusters (15).

Nine T3SS2 effector proteins in *V. parahaemolyticus* have been identified to date (VopA, VopT, VopL, VopV, VopC, VopZ, VPA1380, VopO, and VgpA) (12, 16–23). These effectors subvert several cellular pathways including those controlling actin cytoskeleton dynamics and innate inflammatory responses (reviewed in (24–26)). These effector proteins are classified as either core or accessory T3SS2 effector proteins based on their distribution among the T3SS2 phylotypes (13). Notably, the presence of multiple uncharacterized genes in the VPaI-7 region raises the possibility that there are additional T3SS2 effector proteins yet to be identified.

In this study, we found that VPA1328, an ORF in the *V. parahaemolyticus* VPaI-7, encodes a novel T3SS2 effector protein. VPA1328, re-named here VopG, due to its similarity to the uncharacterized *V. cholerae* effector VopG, is secreted in a T3SS2-dependent fashion. Comparative genomic and phylogenetic analyses revealed that VPA1328 and VopG are members of a larger family of T3SS2 effector proteins encoded within the T3SS2 clusters of vibrios outside of *V. parahaemolyticus* and *V. cholerae* including in *V. mimicus* and *V. diabolicus* and the marine bacterium *Shewanella baltica*. The association of *vopG* genes with insertion sequence elements in several of these clusters suggests independent horizontal gene transfer or rearrangement events in these loci. Furthermore, VopG proteins have a conserved domain that exhibits sequence and predicted structural similarity to the serine/threonine kinase domain in the well-characterized NleH family of T3SS effector proteins. These effectors have been linked to subversion of host cell survival pathways and suppression of innate immunity in infected cells (27–29). However, VopG did not block host cell death or IL-8 secretion in tissue culture-based infections, suggesting a yet undefined role for VopG during infection or functional redundancy with other T3SS2 effector proteins.

## RESULTS

### VPA1328 is a VopG homolog that is secreted and translocated into host cells by the *Vibrio parahaemolyticus* T3SS2

We carried out BLASTp-based homology searches of the *V. parahaemolyticus* VPaI-7 genomic island as a way to identify candidate new T3SS2 effector proteins. This approach suggested that VPA1328 (VP_RS21530 in the latest genome annotation of strain RIMD2210633) is a putative T3SS2 effector protein (**Table S2**). VPA1328 is predicted to encode a 260 amino acid protein that shares ~42% amino acid sequence identity with the T3SS effector protein VopG, encoded in the phylogenetically related T3SS2 in *V. cholerae* AM-19226 (30) (**Figure 1A and 1B**). The function of VopG remains unknown, but it is secreted and translocated by the *V. cholerae* T3SS2 and contributes to host cell cytotoxicity and colonization in a mouse model of infection (30) suggesting an important role for this effector in virulence (30). Even though VPA1328 and VopG are located in different locations within their respective T3SS2 clusters, their sequence similarity and presence in phylogenetically related T3SSs suggests that VPA1328 is a VopG homolog that functions as a *V. parahaemolyticus* T3SS2 effector protein. Below we refer to VPA1328 as VopG.

**Figure 1.**
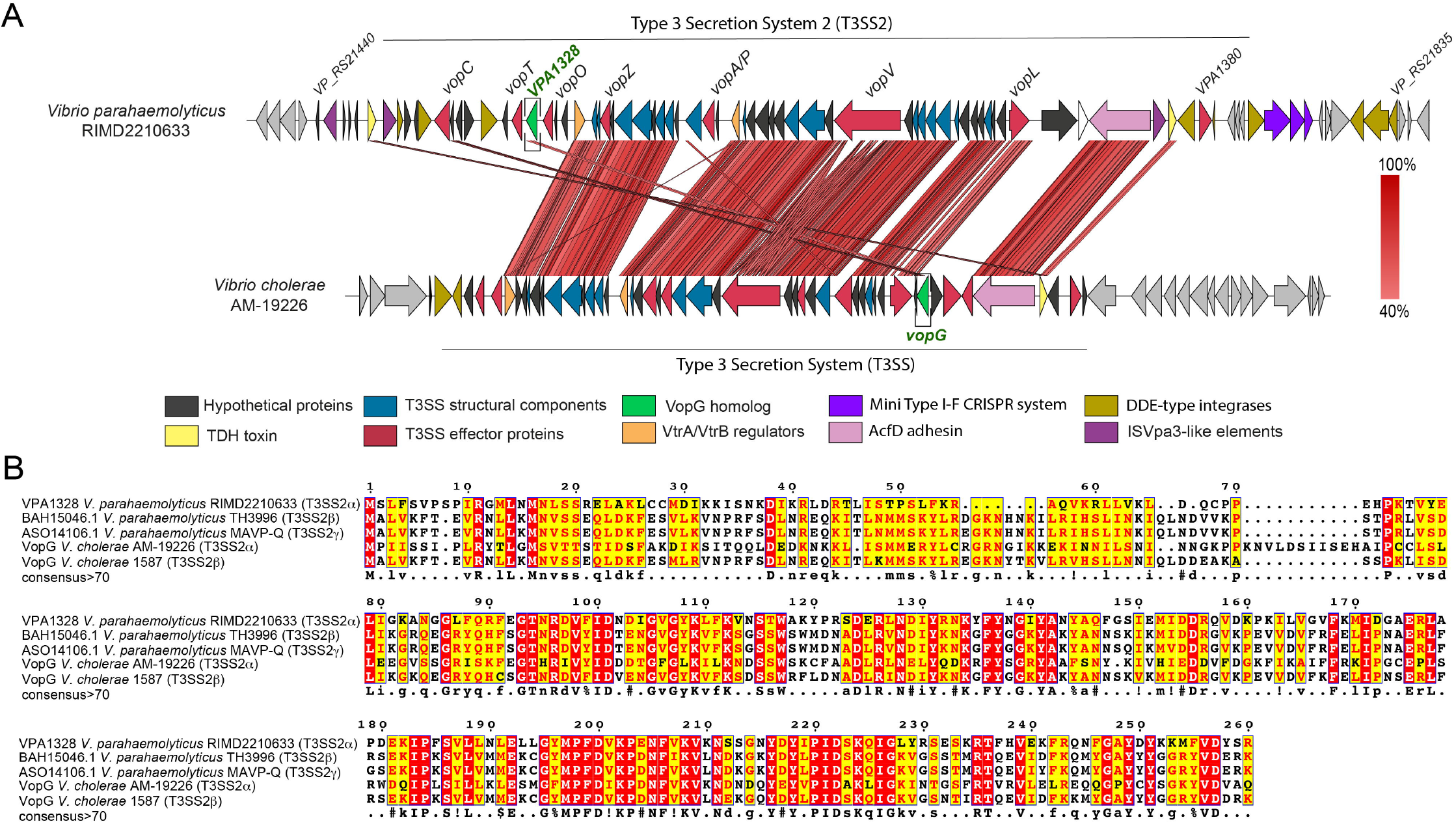
VPA1328, an ORF in the *V. parahaemolyticus* RIMD2210633 T3SS2 gene cluster, is similar to the VopG effector protein encoded within the *V. cholerae* AM-19226 T3SS2 gene cluster. (A) Schematic depiction of a comparison of the T3SS2 gene clusters in *V. parahaemolyticus* RIMD2210633 and *V. cholerae* AM-19226. BLASTn alignment was performed and visualized using EasyFigure (B) Multiple Sequence alignment of VPA1328 and VopG homologs. BLASTp alignments were performed using T-Coffee Expresso and visualized by ESPript 3.0. Amino acids with a red background correspond to positions with 100% identity, amino acids with a yellow background correspond to positions with >70% identity.

Next, we tested if VopG (VPA1328) is secreted and whether its secretion requires the *V. parahaemolyticus* T3SS2. In these experiments, *V. parahaemolyticus* WT strain RIMD2210633 and isogenic T3SS1, T3SS2, and T3SS1/T3SS2-deficient mutant strains (Δ*vscn1,* Δ*vscn2* and Δ*vscn1*Δ*vscn2*, respectively) were grown under conditions (27) that induce expression of T3SS2 (LB 0.04% bile) (31). To detect VopG, these strains were transformed with pVPA1328-CyaA, a plasmid that harbors a translational fusion between the VPA1328 ORF (VopG) and the adenylate cyclase domain (CyaA) of plasmid pCyaA. This construct enables immunoblot detection of VopG in cell lysates and culture supernatants using anti-CyaA antibodies. A VopV-CyaA fusion (pVopV-CyaA) was included as a positive control for T3SS2-dependent secretion (19).

A band corresponding to the predicted size of the VopG-CyaA fusion (~74kDa, along with some lower molecular weight species likely corresponding to degradation products) was observed in cell lysates from the WT strain harboring pVopG-CyaA, but not a control strain harboring the empty vector pCyaA (**Figure 2A**). VopG was only detected in supernatants when the WT (pVopG-CyaA) strain was grown under T3SS2 inducing (LB 0.04% bile) and not in culture supernatants in strains lacking a functional T3SS2 (**Figure 2A**), suggesting that its secretion requires T3SS2 activity. Interestingly, previous transcriptomic analysis showed that expression of VPA1328 was increased by the presence of bile and controlled by VtrB, the master regulator of T3SS2 expression, suggesting that it is part of the VtrB regulon (32).

**Figure 2.**
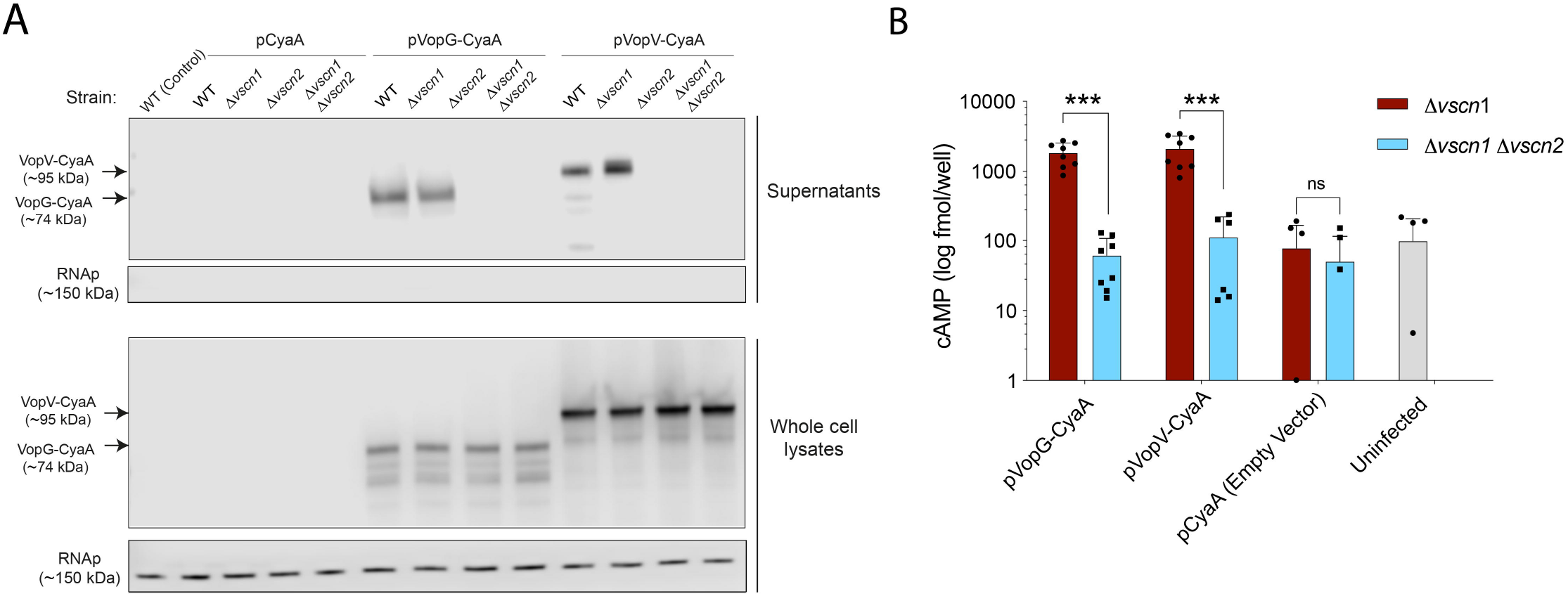
VPA1328 is secreted and translocated in a T3SS2 dependent fashion. (A) Immunoblots of culture supernatants and whole cell lysates using anti-CyaA antibodies for CyaA tagged VPA1328 (VopG) and VPA1357 (VopV) in WT (A) and isogenic Δ*vscn1,* Δ*vscn2* and Δ*vscn1*Δ*vscn2* mutant strains. Immunoblots for RpoB were used as a control for cell lysis. (B) Translocation of VopG-CyaA and VopV-CyaA fusions was assessed via determination of intracellular cAMP levels of infected Caco-2 cells. Error bars indicate standard deviations from 2 independent biological replicates each with 4 technical replicates. Asterisks indicate significant differences (t-test, *** P<0.001).

Analyses of VopG secretion from Δ*vscn1* (T3SS1-deficient) and Δ*vscn2* (T3SS2-deficient) strains strongly support the idea that VopG secretion requires T3SS2 and not T3SS1. When secretion by T3SS1 or T3SS2 or both T3SS was disabled by deletion of their respective ATPases, there was similar expression of VopG in cell lysates (**Figure 2A**); however, VopG was only detected in supernatants from the strain where T3SS1 was inactivated but not when T3SS2 was inactive. An identical pattern was observed with VopV, a known T3SS2 substrate (**Figure 2A**). The cytosolic RNA polymerase beta subunit (RpoB) was not detected in any of the culture supernatant samples, indicating that detection of VPA1328 in culture supernatants was not a consequence of bacterial lysis.

We also tested whether VPA1328 (VopG) is translocated into infected host cells by *V. parahaemolyticus’ T3SS2*. Caco-2 cells were infected with *V. parahaemolyticus* strains harboring the effector-CyaA reporter fusions described above, and the amount of intracellular cAMP generated by each translocated effector was measured using an enzyme-linked immunosorbent assay (ELISA). As shown in **Figure 2B**, after 1 hour of infection, cAMP levels generated by VopV-CyaA (positive control) and VopG-CyaA were similar and significantly higher in cells infected with *V. parahaemolyticus* strains harboring a functional T3SS2 (Δ*vscn1*, T3SS2+) than in cells infected with a T3SS deficient strain (Δ*vscn1* Δ*vscn2,* T3SS-), which exhibited background cAMP levels (**Figure 2B**). Together, these observations demonstrate that VopG is secreted and translocated into host cells in a T3SS2-dependent fashion and given its similarity to the *V. cholerae* VopG effector, strongly support the notion that VopG is a novel *V. parahaemolyticus* T3SS2 effector protein.

### VopG homologs are widely distributed in Vibrios harboring T3SS2 clusters

The presence of a VopG homolog encoded within the T3SS2 gene cluster in *V. parahaemolyticus* RIMD2210633 prompted us to investigate if additional VopG homologs are present among distinct T3SS2 phylotypes. The VPA1328 sequence was used as a query to identify potential VopG homologs by sequential BLASTn, BLASTp and tBLASTx searches, using publicly available bacterial genome sequences. With cut-off values of 60% sequence coverage and 40% sequence identity, 2044 candidate VopG homologs were identified, including 123 non-redundant protein sequences (Figure 3A, 3B and Table S3). The majority of the VopG homologs (86%, n=1764) were encoded in *V. parahaemolyticus* strains and in *V. cholerae* strains (12.5%, n=256), but homologs were also identified in *V. mimicus* (0.2%, n=5), *V. diabolicus* (0.04%, n=1), *Vibrio sp* (0.5%, n=11) and in *Shewanella* strains (0.3%, n=7); i.e., in most species known to harbor T3SS2 gene clusters. Interestingly, a T3SS2 gene cluster was not previously identified in *V. diabolicus*, a marine organism. However, it is important to note that not all *Vibrio* species, e.g., *Vibrio anguillarum* (33) which harbor T3SS2 gene clusters, encode VopG homologs. Thus, even though VopG is widely distributed, this putative effector protein is not a universal component of the T3SS2.

**Figure 3.**
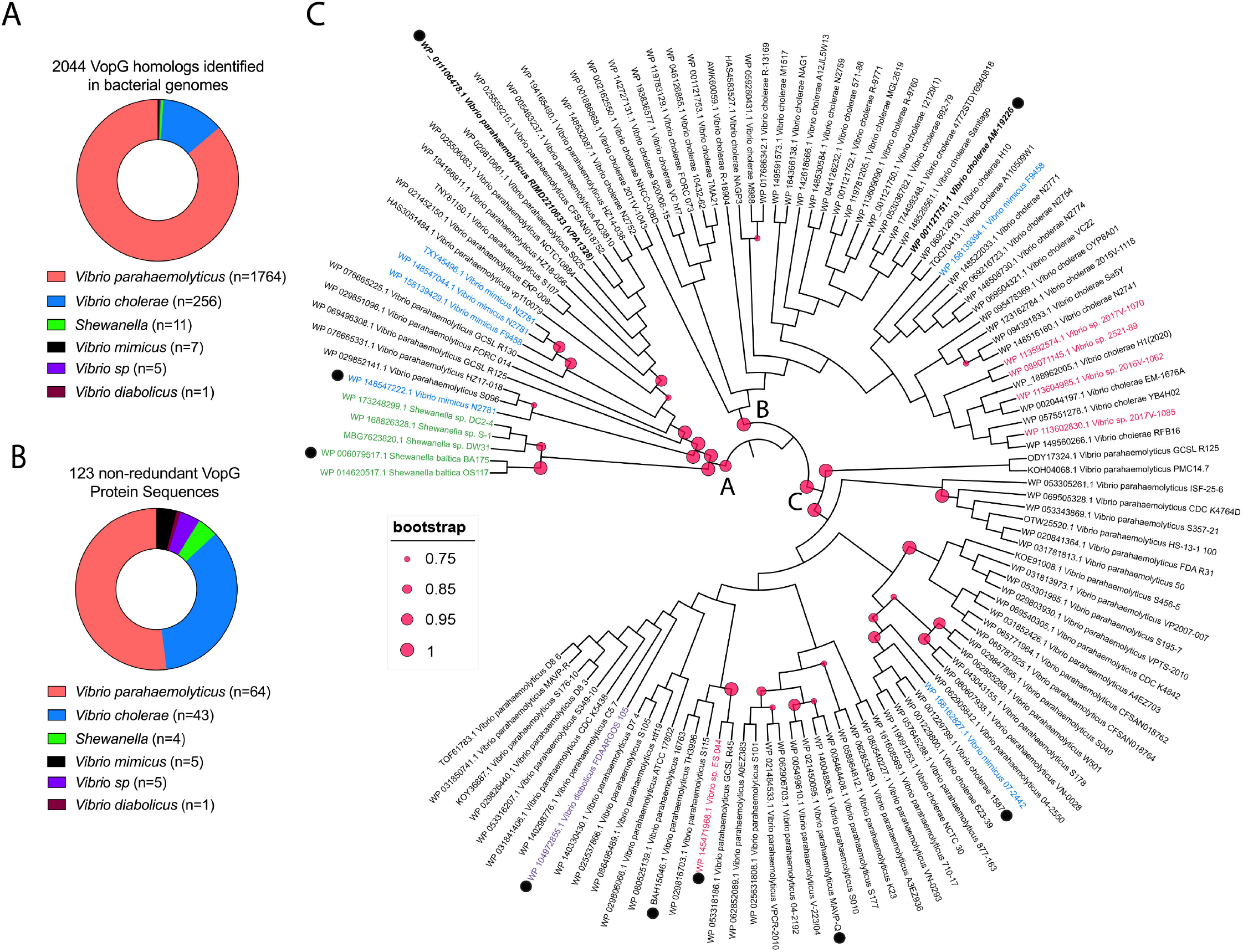
VopG homologs are widely distributed among *Vibrio* and *Shewanella* species. (A) Number of *Vibrio* sp and *Shewanella* sp isolates where VopG homologs were identified. (B) Distribution of the 123 non-redundant VopG protein sequences in different *Vibrio* and *Shewanella* species. (C) Phylogenetic analysis of the 123 non-redundant VopG homologs identified in this study. Phylogenetic analysis was performed with MEGA and visualized by iTOL. Distinct bacterial species are highlighted in different colors. VopG homologs used in the multiple sequence alignment of Fig 5A are highlighted with a black dot.

Next, we evaluated the sequence relatedness of VopG homologs using phylogenetic analysis of the 123 non-redundant VopG sequences. Three distinct clades (A, B and C) of VopG proteins were identified **(Figure 3C)** but no clear correlation was found between these clades and T3SS2 phylotypes. For example, the VopG homologs of *V. cholerae* strain AM-19226 and *V. parahaemolyticus* RIMD2210633 (VPA1328) clustered in different clades (B and A, respectively) despite the fact that both these T3SS2 belong to the T3SS2α phylotype. The lack of correlation between the VopG clades and T3SS2 phylotypes suggests that VopG effectors have to some extent evolved independently of the T3SS2 machinery that delivers them to host cells.

Comparative genomic analyses were carried out to gain insights into variation of the genomic contexts of *vopG* genes within different T3SS2 gene clusters. Genome sequences from representatives of each clade of the VopG phylogenetic tree, including at least one genome for each different *Vibrio* species were used for these comparisons. As shown in **Figure 4**, the overall genetic structure of these T3SS2 gene clusters is highly conserved, particularly in the regions encoding structural components of the T3SS2 apparatus. In most T3SS2 gene clusters, the relative position of *vopG* was similar with the exception of *V. parahaemolyticus* RIMD2210633. However, the nucleotide sequences and ORFs that are adjacent to the *vopG* homologs differed in most of the 7 clusters analyzed in **Figure 4**. In several of these cases, *vopG* was found to be close to sequences related to insertion sequence (IS) elements. This association raises the possibility that IS elements can promote the mobility of *vopG* loci and potentially account for the variations in the genetic contexts of these loci within different T3SS2 gene clusters.

**Figure 4.**
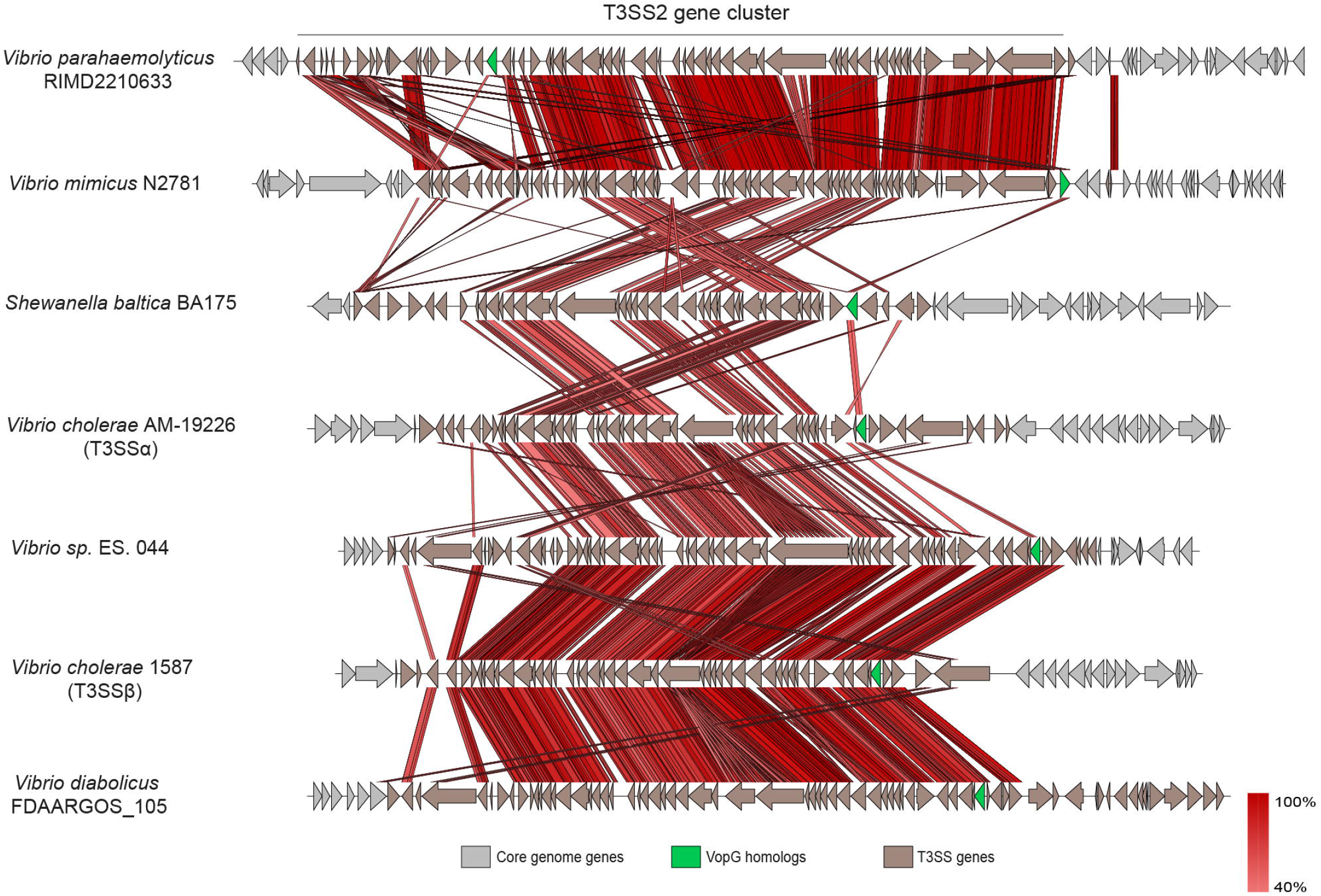
Location of *vopG* homologs within T3SS2 gene clusters. BLASTn alignments were performed and visualized using EasyFigure

Consistent with this idea, we identified two *vopG* homologs (FORC14_RS05860 and FORC14_RS06170) encoded in the T3SS2 gene cluster of *V. parahaemolyticus* strain FORC014. Analysis of their respective genetic contexts revealed that one of these *vopG* genes (FORC14_RS05860) is located at the end of the T3SS gene cluster and is flanked by IS200-like mobile genetic elements (**Figure S1A)**. These elements have high sequence identity to the ISVpa3 insertion sequence. ISVpa3 is an insertion sequence located adjacent to each copy of the TDH gene in *V. parahaemolyticus* strain RIMD2210633 and linked in some strains to deletion of TDH (34). While the T3SS2 gene cluster of *V. parahaemolyticus* RIMD2210633 has 3 copies of these ISVpa3 elements, *V. parahaemolyticus* strain FORC014 has 6 of these elements, 2 of them flanking one of the *vopG* homologs at the end of the cluster (**Figure S1B**). Sequence analysis showed that the 2 *vopG* homologs in strain FORC014 share 69% nucleotide identity (**Figure S2**). Both the sequence divergence of these 2 *vopG* genes and the mobile genetic elements flanking FORC14_RS05860 suggest that this *vopG* homolog was independently acquired, potentially via a horizontal gene transfer event, and not a duplication of FORC14_RS06170.

While the presence of a T3SS2 gene cluster in *Shewanella baltica* species has been inferred due to the presence of the *vscn2* gene in strain *Shewanella baltica* BA175 and *Shewanella baltica* OS183 (35), information regarding the distribution and genetic context of the T3SS2 gene cluster in this genus has not been reported. We found that the *Shewanella* T3SS2 is located within a genomic island inserted between the SBAL678_RS45345 and SBAL678_RS45350 ORFs of reference strain OS678 (**Figure S1B**). This genomic island includes 45 ORFs. Most of these genes encode structural components of the T3SS2 apparatus. Interestingly, not every T3SS2 gene cluster identified in *Shewanella* harbors a VopG-encoding gene (**Figure S1B**).

### VopG proteins have sequence and predicted structural similarity to the NleH family of serine/threonine kinases

The amino acid sequence conservation of the 123 VopG homologs was analyzed and depicted using WebLogo. The analysis showed a particularly striking conservation in the C-termini of these amino acid sequences (**Figure S3**), suggesting that this region of VopG includes a functional domain. To gain clues regarding the function of VopG proteins, we used the structure-based homology tools HHpred (36) and pGenTHREADER (37). For these analyses, the amino acid sequence of *V. parahaemolyticus* VPA1328 was used as a representative of the VopG family of effectors. Both algorithms detected a region in the VopG C-terminus with similarity to NleH effectors, e.g., HHpred analysis uncovered the presence of a region of 22 amino acids in VPA1328 (position 190-212) with identity to the T3SS effector proteins NleH1 from *Escherichia coli* O157:H7 strain Sakai (PDB: 4LRJ_B) and OspG from *Shigella flexneri* 2a strain 301 (PDB: 4Q5E_A).

NleH1 and OspG are members of the NleH of family of T3SS effectors (38–40). These proteins are translocated by the T3SSs of different bacterial species and act as serine/threonine kinases in host cells (39). There are 6 members of this family of effector proteins, including the NleH1 and NleH2 proteins of *E. coli* O157:H7 str. Sakai, OspG from *Shigella flexneri* 2a strain 301, NleH of *Citrobacter rodentium* strain DBS100, SboH of *Salmonella bongori* NCTC 12419 and YspK of *Yersinia enterocolitica* strain 8081 (27, 29, 41–44). The serine/threonine kinase domain of these effectors is distantly related to eukaryotic regulatory kinases; moreover, functional studies have suggested that these effectors can perturb the NF-κB pathway, interfere with innate immune responses and inhibit apoptosis in infected host cells (27–29).

Multiple sequence alignment of representatives of the NleH and VopG family of T3SS effector proteins were carried out to gain further insight into their similarity. Representatives from each clade of VopG proteins were included in these analyses. The analysis showed that the greatest similarity between VopG and NleH proteins is found in their C-termini in the region that includes the characterized NleH serine/threonine kinase domain; in contrast, their N-termini differ in both length and sequence (**Figure 5A and Figure S4**). VopG proteins contain all the critical amino acid residues and motifs important for kinase activity, including the conserved catalytic residues glycine of the G-rich loop, the aspartic acid (D) and the asparagine (N) of the catalytic loop (alignment position 200–205 in VPA1328) and the PID motif of the activation loop (alignment position 220–233 in VPA1328). In addition, VopG proteins also share the invariant lysine (alignment position 109) involved in the autophosphorylation of the NleH family, and which has been used as a proxy to measure kinase activity (28) (**Figure 5A and 5B**). Thus, VopG family effector proteins harbor a NleH-like C-terminal serine/threonine kinase domain. Phylogenetic analysis of bacterial serine/threonine kinase domains also revealed the similarity of the kinase domains of VopG and NleH proteins (**Figure 6A**). The VopG proteins clustered closer to the NleH proteins on this tree than to non-NleH serine threonine kinases from *Legionella pneumophila, Yersinia pestis*, and *Salmonella enterica*.

**Figure 5.**
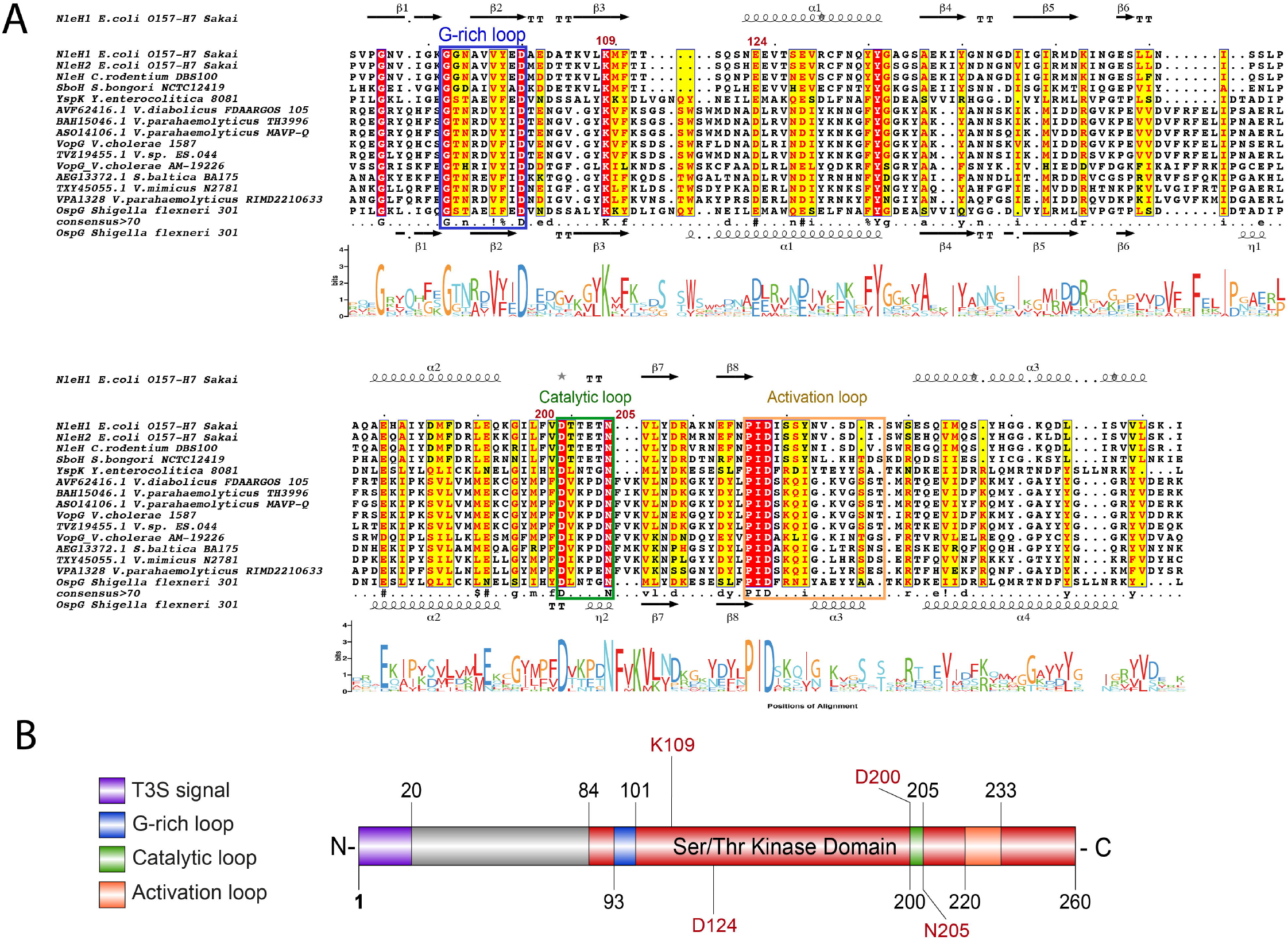
Conservation of the C-terminal domains of VopG homologs. (A) Multiple sequence alignment and Weblogo analysis of the C-terminal domains of VopG proteins (aminoacids 84-260 in VPA1328) with the serine/threonine kinase domains of NleH proteins. G-rich loop, catalytic and activation loops of the kinase are highlighted in colored boxes matching the schematic diagram in (B). BLASTp alignment was performed using T-Coffee Expresso and visualized by ESPript 3.0. Amino acids within a red background correspond to positions with 100% identity, amino acids with a yellow background correspond to positions with >70% identity. The secondary structure of NleH1 and OspG is shown flanking the alignment (α, alpha helices; β, beta sheets; T, turns). (B) Schematic representation of VPA1328 highlighting the presence of the T3S signal as well as conserved regions and key catalytic residues of the putative serine/threonine kinase domain identified in (A).

**Figure 6.**
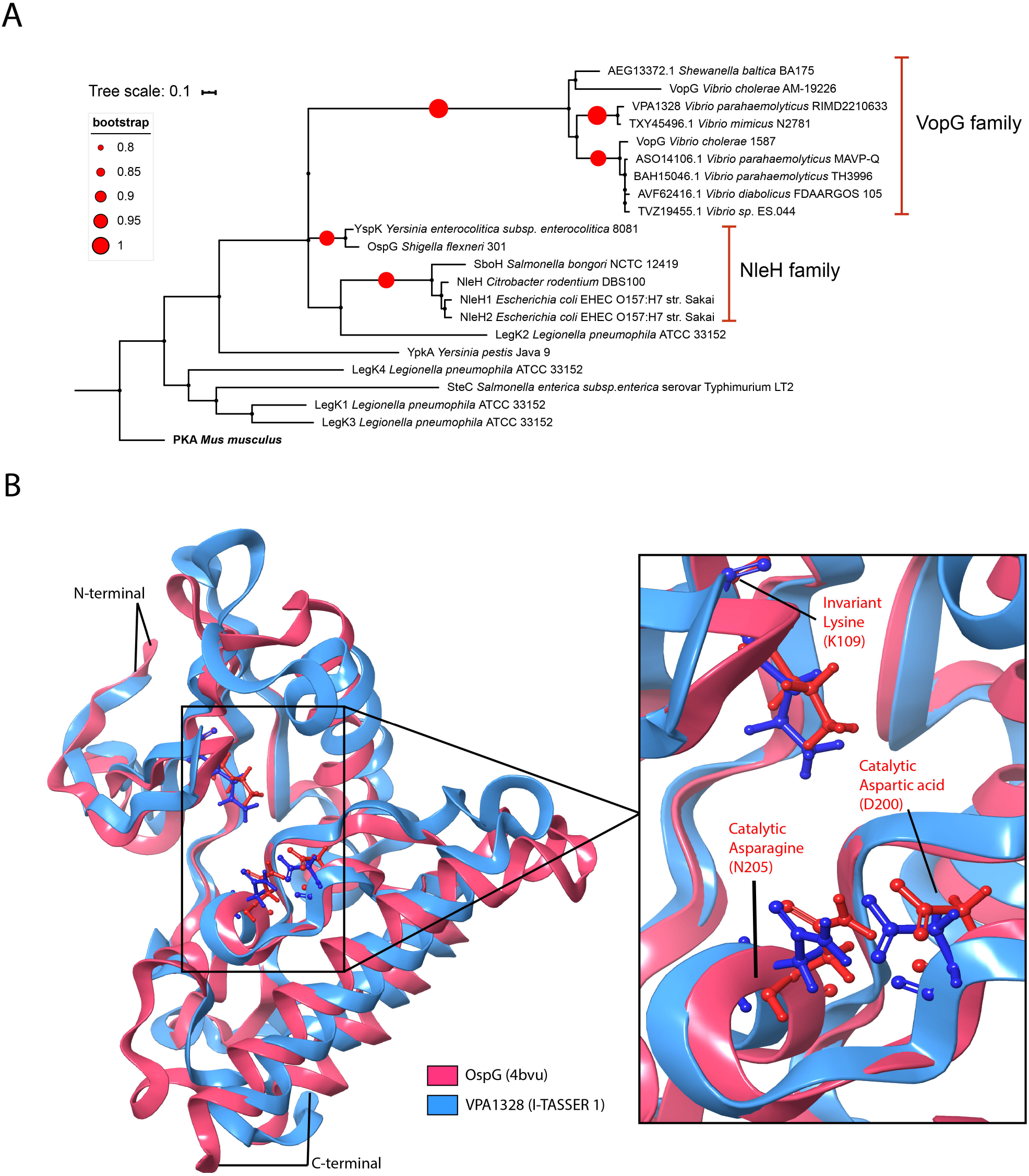
VopG homologs contain a NleH-like serine/threonine kinase domain. (A) Phylogenetic analysis of bacterial serine/threonine kinases. The analysis was performed with MEGA and visualized by iTOL. (B) Comparative homology model of the C-terminal domain of VPA1328 (aminoacids 84-260) superimposed on the known structure OspG (PDB: 4BVU). The inset depicts the catalytic domain of OspG and the superimposed predicted structure of this region in VPA1328. Homology modelling was performed using the I-TASSER pipeline and visualized with MAESTRO.

We derived a 3D structural model of the C-terminal domain of VPA1328 using comparative homology modeling with I-TASSER (45) to gain further insight into the serine/threonine kinase domain of VopG proteins. In accord with the HHPred and pGenTHREADER analyses, I-TASSER identified NleH1, NleH2 and OspG as suitable models for comparative homology models using the crystal structures available for these proteins. Five models were obtained, and I-TASSER model 1 was chosen based on its error estimation, TM-score and RMSD values (**Figure S5**). As shown in **Figure 6B**, this model revealed the remarkable similarity of the predicted structure of the VPA1328 kinase domain with the NleH kinase domain. The structure of the catalytic pocket, including the positions of the predicted catalytic amino acid side chains (K109, D201 and N205) in VPA1328 and OspG structure overlap **(Figure 6B)**, strongly supporting the notion that the VopG family of proteins encode a NleH-like serine/threonine kinase domain.

### VopG does not modulate T3SS2-mediated cytotoxicity or inhibit IL-8 production

NleH family effector proteins inhibit IL-8 expression (27) and host cell death during infection (27–29). Since *V. parahaemolyticus’* capacity to suppress IL-8 secretion and to induce host cell death is partially dependent on a functional T3SS2 (19, 46), we evaluated if VopG contributes to these processes. A *V. parahaemolyticus* mutant strain harboring a deletion of *vopG* in the Δ*vscn1* genetic background was constructed to assess if *vopG* modulates T3SS2-dependent killing of intestinal Caco-2 cells. As expected, Caco-2 cells were killed (<50% survival within 3.5 hours of infection) by a *V. parahaemolyticus* strain harboring a functional T3SS2 (Δ*vscn1*, T3SS2+), whereas cells infected with a *V. parahaemolyticus* strain lacking both T3SSs (Δ*vscn1,* Δ*vscn2*) were not (Figure 7A). However, the absence of *vopG* (Δ*vscn1* Δ*vopG*, T3SS2+) did not alter survival of host cells infected with a functional T3SS2 (overlap of red and orange survival curves in Figure 7A), suggesting that *vopG* does not modulate T3SS2-dependent cytotoxicity.

**Figure 7.**
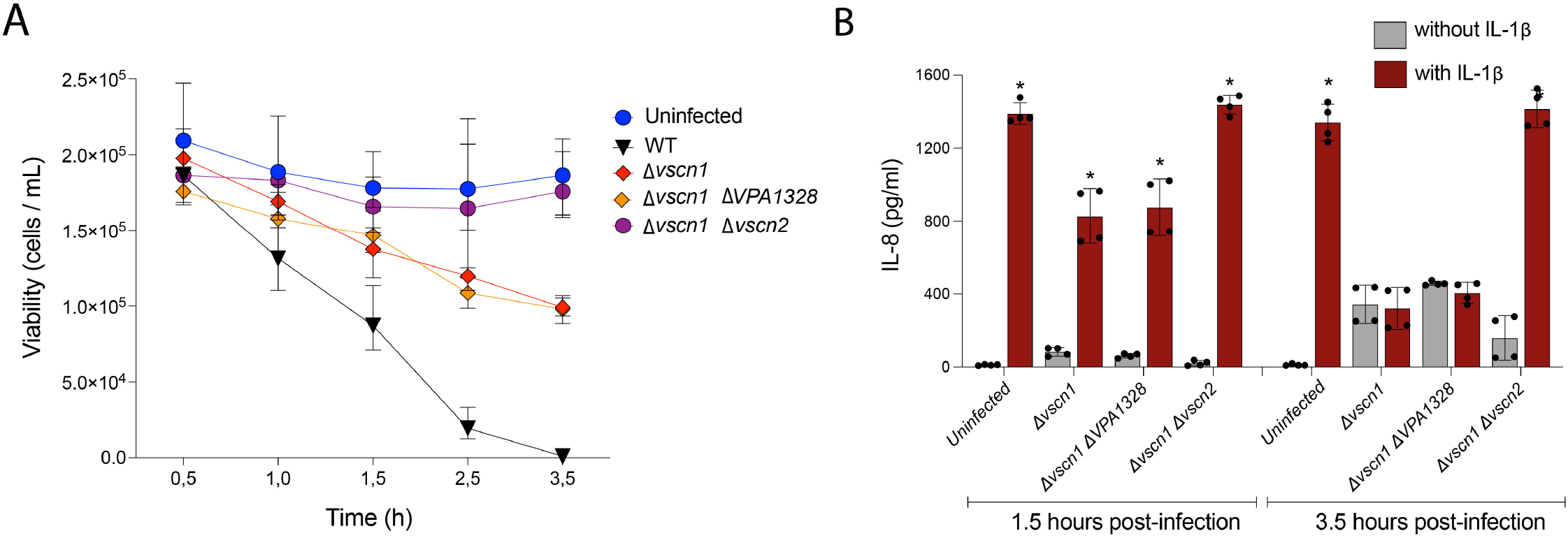
VopG does not modulate T3SS2-dependent cytotoxicity or suppression of IL-8 production. (A) Survival kinetics of Caco-2 cells infected with WT *V. parahaemolyticus* or derivatives with inactive T3SS1 (Δ*vscn1*), inactive T3SS1 without *vopG* (Δ*vscn1,* ΔvopG) or inactive T3SS1 and T3SS2 (Δ*vscn1,* Δ*vscn2*). Cell viability was evaluated by trypan blue exclusion tests at the indicated times. (B) Levels of IL-8 secretion (pg/mL) from HeLa cells infected with the indicated *V. parahaemolyticus* strains in the presence or absence of IL-1β (25 ng/mL) as an inducer;. Error bars indicate standard deviations from 2 biological replicates each with 2 technical replicates. Asterisks indicate significant differences (multiple paired t-test, P<0.001) in comparison to the respective infected cells without IL-1β induction.

We then tested whether VopG contributes to T3SS2-dependent suppression of IL-8 production in infected cells. HeLa cells were infected with *V. parahaemolyticus* strains for 1.5 or 3.5 hours and then IL-8 secretion was stimulated by incubating the cells with IL-1β for 90 minutes, as previously described (19), Infection by *V. parahaemolyticus* inhibited IL-8 secretion in a T3SS2-dependent fashion, but the absence of *vopG* did not influence this phenotype (**Figure 7B**). Together, these data suggest that VopG does not modulate T3SS2-dependent host cell death or inhibit IL-8 production in infected cells.

## DISCUSSION

While all *Vibrio parahaemolyticus* strains harbor a T3SS1, a hallmark of the pandemic *Vibrio parahaemolyticus* O3:K6 clone and most human clinical *Vibrio parahaemolyticus* isolates is the presence of a second and phylogenetically distinct T3SS2. The latter T3SS is essential for both intestinal colonization and virulence in some animal models of disease (10, 12). Here, we found that a T3SS2 ORF (VPA1328) likely corresponds to a novel *V. parahaemolyticus* T3SS2 effector protein. This ORF, which is secreted in a T3SS2-dependent fashion, bears similarity to the VopG effector found in the *V. cholerae* AM-19226 T3SS2. The function of the latter VopG protein is unknown, but it has been shown to be translocated to host cells and linked to *V. cholerae* AM-19226’s pathogenicity. Bioinformatic analyses uncovered 123 non-redundant VopG-like proteins encoded in all 3 phylotypes of T3SS2 clusters in diverse *Vibrio* species and in *S. baltica*. Interestingly, the evolutionary history of the T3SS2 phylotypes does not appear to correspond with the evolution of the 3 clades of VopG proteins that were uncovered by phylogenetic analysis. We found that the highly conserved C-terminal domains of VopG proteins bear striking structural similarity to the serine/threonine kinase domain of the NleH family of effectors found in enteric pathogens such as EHEC and *Shigella* (OspG). Thus, our findings support the idea that VopG effectors function as serine/threonine kinases in host cells.

The *V. cholerae* AM-19226 effector VopG had been classified as a *V. cholerae* specific T3SS effector (30, 47) but our analyses showed that VopG homologs belong to a larger family of putative effector proteins that is widely distributed among *Vibrio* species including *V. parahaemolyticus*, *V. cholerae*, *V. mimicus* and *V. diabolicus* as well as in strains of *Shewanella baltica*. Recently, Matsuda et al proposed to classify T3SS2 effectors proteins as “core” effectors if they are conserved in both *V. parahaemolyticus* and non-O1/non-O139 *V. cholerae* (13) and as “accessory” effectors (13) if they are not. According to this classification, our work suggests that VopG corresponds to a core effector protein due to its presence in multiple *Vibrio* species. However, VopG homologs are not present in the T3SS2 gene clusters identified in all *Vibrio* species; e.g, the T3SS2 cluster in *Vibrio anguillarum* (33) lack a VopG homolog and not all clusters in *V*. *mimicus* (48) encode a recognizable VopG.

T3SS2 gene clusters are classified into three phylotypes (T3SS2α, T3SS2β and T3SS2γ) that are believed to have been acquired through horizontal gene transfer events (13, 15, 48). Even though VopG homologs are not universally found in all T3SS2 gene clusters, we identified VopG homologs in all three T3SS2 phylotypes. Phylogenetic analysis identified three distinct VopG clades (**Figure 3C**). These VopG clades did not correlate with T3SS2 phylotypes, i.e., all three clades were found in each T3SS2 phylotype. The apparent independent evolution of T3SS2 phylotypes and VopG clades supports the possibility that *vopG* genes have been independently acquired by different T3SS2 lineages.

The absence of *vopG* genes from certain T3SS2 clusters could be explained either by loss of *vopG* loci due to deletion event(s) or independent acquisition of *vopG* in some T3SS2 clusters. The presence of a second *vopG* homolog flanked by IS elements in *V. parahaemolyticus* strain FORC014 suggests insertion sequences may play a role in mobilizing *vopG* genes. These sequences bear similarity to the ISVpa3 insertion sequence first described in *V. parahaemolyticus* RIMD2210633 (34). Since insertion sequences have been shown to shape bacterial genomic islands through rearrangements, insertion, and deletion events, it is plausible that ISVpa3-like elements have shaped the evolution of T3SS2 gene clusters through similar mechanisms. The apparent mobility of *vopG* loci adds an additional layer of complexity to our understanding of T3SS2 clusters. That is, these clusters appear to have been spread via horizontal gene transfer events among marine bacteria and their repertoire of effector proteins appears to be “tunable” through independent horizontal gene transfer or rearrangement events.

Analysis of the amino acid sequences of the 123 non-redundant VopG homologs identified here revealed a particularly high degree of conservation in their C-termini. This region of VopG proteins was found to be very similar to the conserved serine/threonine kinase domain in the NleH family of T3SS effector proteins. Thus, the conservation of this part of VopG effectors is likely explained by the presence of a functional kinase domain. Structural predictions, which showed that VopG proteins contain all the residues that constitute that catalytic pocket of NleH proteins, strongly support this hypothesis.

The NleH family of effectors contain a eukaryotic-like serine/threonine kinase domain that independently evolved in bacteria (38, 39). VopG homologs harbor each of the key residues described in the NleH family of protein kinases. The classification of the NleH proteins as a distinct bacterial kinase family was made through structure-based phylogenetic analysis (38). Phylogenetic analysis showed that the C-termini of VopG proteins have more similarity to the kinase domain of NleH proteins than to other bacterial protein kinases (**Figure 6A**), but further structural information is required to determine if VopG proteins are novel members of the NleH family or represent a distinct family on their own.

Although the conservation of key catalytic residues in VopG and NleH proteins provides evidence that supports the notion that VopG proteins are functional serine/threonine kinases, it is more problematic to speculate that their biological function is conserved as well. To date, NleH proteins have been shown to perturb the NF-κB pathway and impact cell survival and innate immune responses during infection through different molecular mechanisms (27–29, 41, 49). Both NleH1 and NleH2 proteins bind the host protein RPS3 leading to inhibition or activation of the NF-κB pathway respectively (27, 50), but only NleH1 suppresses IL-8 expression during EHEC infection (27). Despite these differences in control of IL-8 expression, both NleH1 and NleH2 inhibit apoptosis through interaction with the Bax Inhibitor 1 protein (28). The *Shigella* OspG protein inhibits the NF-κB pathway by inhibiting the proteasomal destruction of IκBα (43) and the SboH protein of *Salmonella bongori* blocks intrinsic apoptotic pathways (29). The *Vibrio parahaemolyticus* T3SS2 causes cell death (18, 51, 52) and suppresses IL-8 secretion through perturbation of the NF-κB pathway (19, 46). While the exact mechanism of T3SS2-mediated host cell death is unknown, the VopZ effector protein plays an important role in inhibiting IL-8 production (19). Our tissue culture-based infection experiments did not reveal that VopG modulates T3SS2-dependent host cell death or IL-8 suppression (**Figure 7**). Two possible scenarios might explain these observations: i) VopG’s contribution to these phenotypes are masked by the redundant effects of additional effectors such as VopZ; or ii) VopG targets host pathways that do not influence cell death or IL-8 synthesis. The N-terminal region of NleH proteins has been linked to substrate recognition and the observed functional differences between the NleH1 and NleH2 proteins (27, 38, 41). In this context, it is tempting to speculate that the sequence divergence observed within the N-terminal region of VopG homologs (**Figure S3 and S4**) has functional implications.

In summary, our work identifies a new family of VopG proteins that are likely T3SS2 effectors. These proteins contain a distinctive NleH-like serine/threonine kinase domain. Future biochemical and structural studies are required to corroborate these predictions. Moreover, defining the role(s) of these effectors in the pathogenicity and/or environmental adaptation of the diverse *Vibrio* and *Shewanella* species that encode them will be fruitful.

## MATERIALS AND METHODS

### Bacterial strains and growth conditions

All bacterial strains and plasmids used in this study are listed in Table S1. *V. parahaemolyticus* RIMD2210633 (3) and its Δ*vscn1*, Δ*vscn2* and Δ*vscn1*Δ*vscn2* derivatives (19) were used in this study. Bacterial strains were routinely cultured in LB medium or on LB agar plates at 37°C. Culture media was supplemented with the following antibiotics and chemicals: 0.04% bovine and ovine bile (Sigma Cat No. B8381); 5μg/mL and 20μg/mL chloramphenicol for *V. parahaemolyticus* and *E. coli* strains, respectively; 1mM IPTG to induce expression vector pCyaA in secretion and translocation assays.

### Eukaryotic cell culture and maintenance conditions

Caco-2 cells (ATCC HTB-37) were maintained in Dulbecco`s Modified Eagle Medium (DMEM) (Gibco) supplemented with 10% fetal bovine serum (FBS) (Gibco) at 37°C in 5% CO_2_. Cells were grown at 37°C with 5% CO_2_ and routinely passaged at 70-80% confluency.

### T3SS2 secretion assays

A reporter fusion was constructed between VPA1328 and the CyaA reporter encoded in plasmid pCyaA (a pMMB207 derivative) (53), generating plasmid pVPA1328 (VopG-CyaA), to investigate VPA1328 (VopG) secretion. A VopV-CyaA fusion (pVopV-CyaA) (19) was used as a positive control for T3SS2-dependent secretion, and the empty plasmid pCyaA was used as a negative control. Each plasmid was transformed into *V. parahaemolyticus* by electroporation as previously described (19). Secretion assays were performed by growing each *V. parahaemolyticus* strain for 1.5 h in LB medium supplemented with 0.04% bile. When cultures reached an OD_600nm_ of 0.5-0.6, 1mM IPTG was added to induce expression of the CyaA reporter fusion protein. After 1.5 h of induction, culture supernatants were collected by two centrifugations, one at 5000 RPM for 20 min and a final centrifugation at 13000 RPM for 5 min. The supernatants were then filtered sterilized through a 0.22-μm filter and concentrated 100-fold by repeated centrifugation at 5000 RPM for 30 min with an Amicon Ultra-15 Centrifugal Filter Unit (Millipore) with a 10 kDa molecular weight cut-off and with 2 washes with 15mL of PBS 1X as an exchange buffer. Prior to concentrating the culture supernatant, BSA (1□mg/mL) was added to serve as a concentration/loading control. Whole-cell lysates were prepared by solubilizing the bacterial pellets in 1X Laemmli buffer. Lysate and supernatant samples were processed for SDS-PAGE analysis by mixing them with loading buffer, boiling for 5 min; they were then and run on 4-20% Mini-PROTEAN^®^ TGX™ Precast Gels (Bio-Rad) per 30 minutes at 200 V. For immunoblot analysis, gels were transferred to iBlot2 transfer stacks of PVDF membranes (Invitrogen) and blocked by EveryBlot Blocking Buffer (Bio-Rad Cat No 12010020). The Pierce Coomassie Plus (Bradford) Assay Kit (Thermo Fisher Cat No. 23236) was used for determination of protein concentrations. Antibodies were used at the following dilutions: anti-CyaA (mouse monoclonal, 1:2,000; Santa Cruz Biotechnology Cat No. sc-13582), anti-RpoB (rabbit monoclonal, 1:2,000; Abcam Cat. No. ab191598), anti-Mouse IgG-HRP (goat polyclonal 1:10,000; Thermo Cat. No. 62-6520) and anti-Rabbit IgG (H+L) Secondary Antibody, HRP (goat polyclonal 1:10,000; Invitrogen Cat. No. G21234). The blots were developed with SuperSignal West Pico Plus substrate (Thermo Fisher Cat No. 35060) and imaging was performed on a C-DiGit Blot Scanner (LI-COR Biosciences). All blots are representative of at least 3 biological replicates.

### Translocation of effector-CyaA fusion proteins

CyaA reporter fusion-based protein translocation assays were performed as previously described (Kodama et al., 2007). Briefly, Caco-2 cells were seeded at 1.5×10^4^ cells/well and cultured in DMEM 10% FBS for 2 days in a 96-well plate at 37°C in 5% CO_2_. *V. parahaemolyticus* RIMD2210633 strains Δ*vscn1* and Δ*vscn1*Δ*vscn2* containing pCyaA, pVopV-CyaA or pVopG-CyaA were grown for 1.5-2 h till OD_600nm_ of 0.6 in LB medium supplemented with 0.04% bile. The infection assay in Caco-2 cells was performed for 1 hr at 37°C, 5% CO_2_ and at multiplicity of infection (MOI) of 50. The intracellular cyclic AMP (cAMP) levels in Caco-2 cells were determined using cAMP Biotrak enzyme immunoassay (EIA) kit (Cytiva Cat No. RPN2251) as described previously (54). Statistical analysis were performed GraphPad Prism version 9 (GraphPad Software, San Diego, California USA, www.graphpad.com).

### T3SS2-dependent cell death and IL-8 secretion assays

For cell survival assays, Caco-2 cells were seeded at 8.0×10^4^ cells/well into 6-well plates and grown for 2 days in complete media. *V. parahaemolyticus* strains were cultured overnight and the next day diluted 1:100 into LB liquid media containing 0.04% bile (to induce T3SS2 expression) and grown for 2 hours until an OD_600nm_ of 0.6. Cells were infected at an MOI=1 and incubated at 37°C with 5% CO_2_. At each time point assayed (0.5, 1.0, 1.5, 2.5, and 3.5 h.), the media was replaced with fresh complete DMEM media supplemented with 100μg/mL of gentamicin. Cells were incubated overnight and surviving cells were quantified either by trypan blue exclusion (0.4% trypan blue) and counted on a haemocytometer (Neubauer cell chamber). For the detection of secreted IL-8, Caco-2 cells were seeded at 1.0×10^5^ cells/well into 12-well plates and cultured in DMEM 10% FBS for 24 hours at 37°C in 5% CO_2_. *V. parahaemolyticus* strains were cultured and the T3SS2 was induced as described above. Cells were infected at an MOI=1 and incubated at 37°C with 5% CO_2_ for 1.5 or 3.5 hours and then, the infection was terminated by addition of gentamicin (100μg/mL). In parallel with gentamicin, the cells were treated with IL-1β (25ng/mL) or left untreated for 90 minutes. IL-8 in culture supernatant was then measured using a human IL-8 ELISA kit (ab46032). Statistical analysis were performed GraphPad Prism version 9 (GraphPad Software, San Diego, California USA, www.graphpad.com).

### Sequence and phylogenetic analysis

Identification of VopG orthologs was carried out using the VPA1328 amino acid and nucleotide sequences as queries in BLASTp, BLASTn, BLASTx, tBLASTn and tBLASTx analyses using publicly available bacterial genome sequences of the NCBI database (December 2020). A 94% sequence length, 40% identity and 60% sequence coverage threshold were used to select positive matches. Sequence conservation was analyzed by multiple sequence alignments using MAFFT (55) and T-Coffee Expresso (56) and visualized by ESPript 3.0 (57). WebLogo analysis was performed using multiple sequence alignments (58). Comparative genomic analysis of the T3SS2 gene clusters was performed using the multiple aligner Mauve (59) and the IslandViewer 4 pipeline (60) and EasyFig v2.2.2. Nucleotide sequences were analyzed by the sequence visualization and annotation tool Artemis version 18.1 (61). Multiple sequence alignments were used for phylogenetic analyses that were performed with the Molecular Evolutionary Genetics Analysis (MEGA) software version 7.0 (62) and visualized by iTOL (63). Phylogenetic trees were built from the alignments by the bootstrap test of phylogeny (2000 replications) using the neighbor-joining (NJ) method with a Jones-Taylor-Thornton (JTT) correction model.

### Remote homology prediction and homology modeling

Remote homology prediction of VPA1328 was performed using HHpred (36) and pGENTHREADER (37) on the PSIPRED server (64). Protein structure models of the VPA1328 C-terminal domain were obtained using I-TASSER (45), a protein structure homology-modelling server. Protein structure visualization and template alignment and superposition were performed using MAESTRO (65).

## ACKNOWLEDGMENTS

We thank members of the Blondel Laboratory for helpful discussions on all aspects of this project and for their comments on the manuscript.

## AUTHOR CONTRIBUTIONS

Nicolas Plaza: Conceptualization, Methodology, Formal analysis, Investigation, Visualization, Data curation, Writing-Original Draft and review and editing.

Italo M Urrutia: Investigation, Visualization, review and editing final manuscript.

Katherine Garcia: Conceptualization, review and editing final manuscript.

Matthew K. Waldor: Conceptualization, Resources, Writing - Review and editing.

Carlos J Blondel: Conceptualization, Methodology, Formal analysis, Investigation, Visualization, Resources, Data curation, Writing - Review and editing, Funding acquisition, Supervision and Project administration.

---

**Figure S1.** Genomic context of *vopG* in *Shewanella baltica* BA175 and *V. parahaemolyticus* FORC014. (A) Schematic of the *vopG* genomic context within T3SS2 gene clusters of *Shewanella* species and within the T3SS gene cluster of *V. parahaemolyticus* FORC014 (B). BLASTn alignment was performed and visualized using EasyFigure

**Figure S2.** Multiple sequence alignment of the DNA sequence of the *vopG* homologs encoded within the T3SS2 gene cluster of *V. parahaemolyticus* strain FORC014. BLASTn alignment was performed using T-Coffee and visualized by ESPript 3.0. Nucleotides within a red background correspond to positions with 100% identity.

**Figure S3.** WebLogo analysis of the multiple sequence alignment of the 123 non-redundant VopG proteins identified in this study.

**Figure S4.** Multiple sequence alignment of the amino acid sequences of the VopG homologs from *V. parahaemolyticus* RIMD2210633 and *V. cholerae* AM-19226 with the NleH1, NleH2 proteins of *E. coli* O157:H7 and OspG protein of *S. sonnei* strain Ss046. BLASTp alignment was performed using M-Coffee and visualized by ESPript 3.0. Amino acids within a red background corresponds to positions with 100% identity and amino acids in a yellow background correspond to positions with over 70% of identity.

**Figure S5.** Estimated accuracy of the five protein structures of the C-terminal (amino acids 84-260) domain of VPA1328 obtained through comparative homology modelling using the I-TASSER pipeline. The locations of the predicted G-rich loop, catalytic and activation loop are highlighted.

**Table S1.** Bacterial strains and plasmids used in this study.

**Table S2.** Analysis of the VPA1328 ORF using different T3SS effector prediction software.

**Table S3.** List of the 2044 total VopG protein sequences and the 123 non-redundant VopG protein sequences identified in bacterial genome databases.

## REFERENCES

1. Letchumanan V, Chan K-G, Lee L-H. 2014. Vibrio parahaemolyticus: a review on the pathogenesis, prevalence, and advance molecular identification techniques. Frontiers in microbiology 5:705.

2. Velazquez-Roman J, León-Sicairos N, Hernández-Díaz L de J, Canizalez-Roman A. 2013. Pandemic Vibrio parahaemolyticus O3:K6 on the American continent. Frontiers in cellular and infection microbiology 3:110.

3. Makino K, Oshima K, Kurokawa K, Yokoyama K, Uda T, Tagomori K, Iijima Y, Najima M, Nakano M, Yamashita A, Kubota Y, Kimura S, Yasunaga T, Honda T, Shinagawa H, Hattori M, Iida T. 2003. Genome sequence of Vibrio parahaemolyticus: a pathogenic mechanism distinct from that of V cholerae. Lancet 361:743–749.

4. Hazen TH, Lafon PC, Garrett NM, Lowe TM, Silberger DJ, Rowe LA, Frace M, Parsons MB, Bopp CA, Rasko DA, Sobecky PA. 2015. Insights into the environmental reservoir of pathogenic Vibrio parahaemolyticus using comparative genomics. Frontiers in microbiology 6:204.

5. Abby SS, Rocha EPC. 2012. The non-flagellar type III secretion system evolved from the bacterial flagellum and diversified into host-cell adapted systems. PLoS genetics 8:e1002983.

6. Park K-S, Ono T, Rokuda M, Jang M-H, Okada K, Iida T, Honda T. 2004. Functional characterization of two type III secretion systems of Vibrio parahaemolyticus. Infection and immunity 72:6659–6665.

7. Portaliou AG, Tsolis KC, Loos MS, Zorzini V, Economou A. 2016. Type III Secretion: Building and Operating a Remarkable Nanomachine. Trends in biochemical sciences 41:175–189

8. Lara-Tejero M, Galán JE. 2019. Protein Secretion in Bacteria. Ecosal Plus 8:245–259.

9. Galán JE, Lara-Tejero M, Marlovits TC, Wagner S. 2014. Bacterial Type III Secretion Systems: Specialized Nanomachines for Protein Delivery into Target Cells. Annu Rev Microbiol 68:1–24.

10. Hubbard TP, Chao MC, Abel S, Blondel CJ, Wiesch PAZ, Zhou X, Davis BM, Waldor MK. 2016. Genetic analysis of Vibrio parahaemolyticus intestinal colonization. Proc National Acad Sci 113:201601718.

11. Piñeyro P, Zhou X, Orfe LH, Friel PJ, Lahmers K, Call DR. 2010. Development of two animal models to study the function of Vibrio parahaemolyticus type III secretion systems. Infection and immunity 78:4551–4559.

12. Ritchie JM, Rui H, Zhou X, Iida T, Kodoma T, Ito S, Davis BM, Bronson RT, Waldor MK. 2012. Inflammation and disintegration of intestinal villi in an experimental model for Vibrio parahaemolyticus-induced diarrhea. PLoS pathogens 8:e1002593.

13. Matsuda S, Hiyoshi H, Tandhavanant S, Kodama T. 2020. Advances on Vibrio parahaemolyticus research in the postgenomic era. Microbiol Immunol 64:167–181.

14. Okada N, Iida T, Park K-S, Goto N, Yasunaga T, Hiyoshi H, Matsuda S, Kodama T, Honda T. 2009. Identification and characterization of a novel type III secretion system in trh-positive Vibrio parahaemolyticus strain TH3996 reveal genetic lineage and diversity of pathogenic machinery beyond the species level. Infection and immunity 77:904–913.

15. Xu F, Gonzalez-Escalona N, Drees KP, Sebra RP, Cooper VS, Jones SH, Whistler CA. 2017. Parallel Evolution of Two Clades of an Atlantic-Endemic Pathogenic Lineage of Vibrio parahaemolyticus by Independent Acquisition of Related Pathogenicity Islands. Appl Environ Microb 83:e01168–17.

16. Hiyoshi H, Okada R, Matsuda S, Gotoh K, Akeda Y, Iida T, Kodama T. 2015. Interaction between the Type III Effector VopO and GEF-H1 Activates the RhoA-ROCK Pathway. Plos Pathog 11:e1004694.

17. Hiyoshi H, Kodama T, Saito K, Gotoh K, Matsuda S, Akeda Y, Honda T, Iida T. 2011. VopV, an F-Actin-Binding Type III Secretion Effector, Is Required for Vibrio parahaemolyticus-Induced Enterotoxicity. Cell Host Microbe 10:401–409.

18. Kodama T, Rokuda M, Park K-S, Cantarelli VV, Matsuda S, Iida T, Honda T. 2007. Identification and characterization of VopT, a novel ADP-ribosyltransferase effector protein secreted via the Vibrio parahaemolyticus type III secretion system 2. Cellular Microbiology 9:2598–2609.

19. Zhou X, Gewurz BE, Ritchie JM, Takasaki K, Greenfeld H, Kieff E, Davis BM, Waldor MK. 2013. A Vibrio parahaemolyticus T3SS effector mediates pathogenesis by independently enabling intestinal colonization and inhibiting TAK1 activation. Cell reports 3:1690–1702.

20. Zhang L, Krachler AM, Broberg CA, Li Y, Mirzaei H, Gilpin CJ, Orth K. 2012. Type III effector VopC mediates invasion for Vibrio species. Cell reports 1:453–460.

21. Trosky JE, Mukherjee S, Burdette DL, Roberts M, McCarter L, Siegel RM, Orth K. 2004. Inhibition of MAPK Signaling Pathways by VopA from Vibrio parahaemolyticus. J Biol Chem 279:51953–51957.

22. Tandhavanant S, Matsuda S, Hiyoshi H, Iida T, Kodama T. 2018. Vibrio parahaemolyticus Senses Intracellular K+ To Translocate Type III Secretion System 2 Effectors Effectively. Mbio 9:e01366–18.

23. Hu M, Zhang Y, Gu D, Chen X, Waldor MK, Zhou X. 2020. Nucleolar c-Myc recruitment by a Vibrio T3SS effector promotes host cell proliferation and bacterial virulence. Embo J e105699.

24. Matsuda S, Hiyoshi H, Tandhavanant S, Kodama T. 2020. Advances on Vibrio parahaemolyticus research in the postgenomic era. Microbiol Immunol 64:167–181.

25. Lingzhi L, Meng H, Dan G, Yang L, Mengdie J. 2019. Molecular mechanisms of Vibrio parahaemolyticus pathogenesis. Microbiol Res 222:43–51.

26. O’Boyle N, Boyd A. 2014. Manipulation of intestinal epithelial cell function by the cell contact-dependent type III secretion systems of Vibrio parahaemolyticus. Frontiers in cellular and infection microbiology 3:114.

27. Gao X, Wan F, Mateo K, Callegari E, Wang D, Deng W, Puente J, Li F, Chaussee MS, Finlay BB, Lenardo MJ, Hardwidge PR. 2009. Bacterial Effector Binding to Ribosomal Protein S3 Subverts NF-κB Function. Plos Pathog 5:e1000708.

28. Hemrajani C, Berger CN, Robinson KS, Marchès O, Mousnier A, Frankel G. 2010. NleH effectors interact with Bax inhibitor-1 to block apoptosis during enteropathogenic Escherichia coli infection. Proc National Acad Sci 107:3129–3134.

29. Fookes M, Schroeder GN, Langridge GC, Blondel CJ, Mammina C, Connor TR, Seth-Smith H, Vernikos GS, Robinson KS, Sanders M, Petty NK, Kingsley RA, Bäumler AJ, Nuccio S-P, Contreras I, Santiviago CA, Maskell D, Barrow P, Humphrey T, Nastasi A, Roberts M, Frankel G, Parkhill J, Dougan G, Thomson NR. 2011. Salmonella bongori provides insights into the evolution of the Salmonellae. PLoS pathogens 7:e1002191.

30. Chaand M, Miller KA, Sofia MK, Schlesener C, Weaver JWA, Sood V, Dziejman M. 2015. Type Three Secretion System Island-Encoded Proteins Required for Colonization by Non-O1/Non-O139 Serogroup Vibrio cholerae. Infect Immun 83:2862–2869.

31. Gotoh K, Kodama T, Hiyoshi H, Izutsu K, Park K-S, Dryselius R, Akeda Y, Honda T, Iida T. 2010. Bile acid-induced virulence gene expression of Vibrio parahaemolyticus reveals a novel therapeutic potential for bile acid sequestrants. PloS one 5:e13365.

32. Livny J, Zhou X, Mandlik A, Hubbard T, Davis BM, Waldor MK. 2014. Comparative RNA-Seq based dissection of the regulatory networks and environmental stimuli underlying Vibrio parahaemolyticus gene expression during infection. Nucleic Acids Res 42:12212–12223.

33. Naka H, Dias GM, Thompson CC, Dubay C, Thompson FL, Crosa JH. 2011. Complete genome sequence of the marine fish pathogen Vibrio anguillarum harboring the pJM1 virulence plasmid and genomic comparison with other virulent strains of V. anguillarum and V. ordalii. Infection and immunity 79:2889–2900.

34. Kamruzzaman M, Bhoopong P, Vuddhakul V, Nishibuchi M. 2008. Detection of a functional insertion sequence responsible for deletion of the thermostable direct hemolysin gene (tdh) in Vibrio parahaemolyticus. Gene 421:67–73.

35. Dong X, Li N, Liu Z, Lv X, Shen Y, Li J, Du G, Wang M, Liu L. 2020. CRISPRi-Guided Multiplexed Fine-Tuning of Metabolic Flux for Enhanced Lacto-N -neotetraose Production in Bacillus subtilis. J Agr Food Chem 68:2477–2484.

36. Zimmermann L, Stephens A, Nam S-Z, Rau D, Kübler J, Lozajic M, Gabler F, Söding J, Lupas AN, Alva V. 2017. A Completely Reimplemented MPI Bioinformatics Toolkit with a New HHpred Server at its Core. J Mol Biol 430:2237–2243.

37. Lobley A, Sadowski MI, Jones DT. 2009. pGenTHREADER and pDomTHREADER: new methods for improved protein fold recognition and superfamily discrimination. Bioinformatics 25:1761–1767.

38. Grishin AM, Cherney M, Anderson DH, Phanse S, Babu M, Cygler M. 2014. NleH Defines a New Family of Bacterial Effector Kinases. Structure 22:250–259.

39. Grishin AM, Beyrakhova KA, Cygler M. 2015. Structural insight into effector proteins of Gram-negative bacterial pathogens that modulate the phosphoproteome of their host. Protein Sci 24:604–620.

40. Grishin A, Cherney M, Condos T, Barber K, Anderson D, Phanse S, Babu M, Shaw G, Cygler M. 2014. Bacterial Effector Kinases. Acta Crystallogr Sect Found Adv 70:C428–C428.

41. Pham TH, Gao X, Tsai K, Olsen R, Wan F, Hardwidge PR. 2012. Functional Differences and Interactions between the Escherichia coli Type III Secretion System Effectors NleH1 and NleH2. Infect Immun 80:2133–2140.

42. García-Angulo VA, Deng W, Thomas NA, Finlay BB, Puente JL. 2008. Regulation of Expression and Secretion of NleH, a New Non-Locus of Enterocyte Effacement-Encoded Effector in Citrobacter rodentiuml11. J Bacteriol 190:2388–2399.

43. Kim DW, Lenzen G, Page A-L, Legrain P, Sansonetti PJ, Parsot C. 2005. The Shigella flexneri effector OspG interferes with innate immune responses by targeting ubiquitin-conjugating enzymes. P Natl Acad Sci Usa 102:14046–14051.

44. Matsumoto H, Young GM. 2006. Proteomic and functional analysis of the suite of Ysp proteins exported by the Ysa type III secretion system of Yersinia enterocolitica Biovar 1B. Mol Microbiol 59:689–706.

45. Yang J, Yan R, Roy A, Xu D, Poisson J, Zhang Y. 2015. The I-TASSER Suite: protein structure and function prediction. Nat Methods 12:7–8.

46. Matlawska-Wasowska K, Finn R, Mustel A, O’Byrne CP, Baird AW, Coffey ET, Boyd A. 2010. The Vibrio parahaemolyticus Type III Secretion Systems manipulate host cell MAPK for critical steps in pathogenesis. Bmc Microbiol 10:329.

47. Alam A, Miller KA, Chaand M, Butler JS, Dziejman M. 2011. Identification of Vibrio cholerae Type III Secretion System Effector Proteins †. Infect Immun 79:1728–1740.

48. Okada N, Matsuda S, Matsuyama J, Park K-S, Reyes C de los, Kogure K, Honda T, Iida T. 2010. Presence of genes for type III secretion system 2 in Vibrio mimicus strains. BMC microbiology 10:302.

49. Grishin AM, Barber KR, Gu R-X, Tieleman DP, Shaw GS, Cygler M. 2018. Regulation of Shigella Effector Kinase OspG through Modulation of its Dynamic Properties. J Mol Biol 430:2096–2112.

50. Pham TH, Gao X, Singh G, Hardwidge PR. 2013. Escherichia coli Virulence Protein NleH1 Interaction with the v-Crk Sarcoma Virus CT10 Oncogene-like Protein (CRKL) Governs NleH1 Inhibition of the Ribosomal Protein S3 (RPS3)/Nuclear Factor κB (NF-κB) Pathway. J Biol Chem 288:34567–34574.

51. Blondel CJ, Park JS, Hubbard TP, Pacheco AR, Kuehl CJ, Walsh MJ, Davis BM, Gewurz BE, Doench JG, Waldor MK. 2016. CRISPR/Cas9 Screens Reveal Requirements for Host Cell Sulfation and Fucosylation in Bacterial Type III Secretion System-Mediated Cytotoxicity. Cell Host Microbe 20:226–237.

52. Hiyoshi H, Kodama T, Iida T, Honda T. 2010. Contribution of Vibrio parahaemolyticus Virulence Factors to Cytotoxicity, Enterotoxicity, and Lethality in Mice l11. Infect Immun 78:1772–1780.

53. Zhou X, Gewurz BE, Ritchie JM, Takasaki K, Greenfeld H, Kieff E, Davis BM, Waldor MK. 2013. A Vibrio parahaemolyticus T3SS Effector Mediates Pathogenesis by Independently Enabling Intestinal Colonization and Inhibiting TAK1 Activation. Cell Reports 3:1690–1702.

54. Zhou X, Ritchie JM, Hiyoshi H, Iida T, Davis BM, Waldor MK, Kodama T. 2012. The hydrophilic translocator for Vibrio parahaemolyticus, T3SS2, is also translocated. Infection and immunity 80:2940–2947.

55. Katoh K, Rozewicki J, Yamada KD. 2017. MAFFT online service: multiple sequence alignment, interactive sequence choice and visualization. Brief Bioinform 20:bbx108-.

56. Notredame C, Higgins DG, Heringa J. 2000. T-Coffee: A Novel Method for Fast and Accurate Multiple Sequence Alignment. JMB 302:205–217.

57. Robert X, Gouet P. 2014. Deciphering key features in protein structures with the new ENDscript server. Nucleic Acids Res 42:W320–W324.

58. Crooks GE, Hon G, Chandonia J-M, Brenner SE. 2004. WebLogo: A Sequence Logo Generator. Genome Res 14:1188–1190.

59. Darling ACE, Mau B, Blattner FR, Perna NT. 2004. Mauve: Multiple Alignment of Conserved Genomic Sequence With Rearrangements. Genome Res 14:1394–1403.

60. Bertelli C, Laird MR, Williams KP, Simon Fraser University Research Computing Group, Lau BY, Hoad G, Winsor GL, Brinkman FS. 2017. IslandViewer 4: expanded prediction of genomic islands for larger-scale datasets. Nucleic Acids Res 45:W30–W35.

61. Carver T, Harris SR, Berriman M, Parkhill J, McQuillan JA. 2012. Artemis: an integrated platform for visualization and analysis of high-throughput sequence-based experimental data. Bioinformatics 28:464–469.

62. Kumar S, Stecher G, Tamura K. 2016. MEGA7: Molecular Evolutionary Genetics Analysis Version 7.0 for Bigger Datasets. Mol Biol Evol 33:1870–1874.

63. Letunic I, Bork P. 2019. Interactive Tree Of Life (iTOL) v4: recent updates and new developments. Nucleic Acids Res 47:gkz239-.

64. Buchan DWA, Jones DT. 2019. The PSIPRED Protein Analysis Workbench: 20 years on. Nucleic Acids Res 47:W402–W407.

65. 2020-4 SR. 2020. MAESTRO. New York.

