## Supplementary figures and images for "Identification of a family of *Vibrio* type III secretion system effectors that contain a conserved serine/threonine kinase domain"

### Figure S1

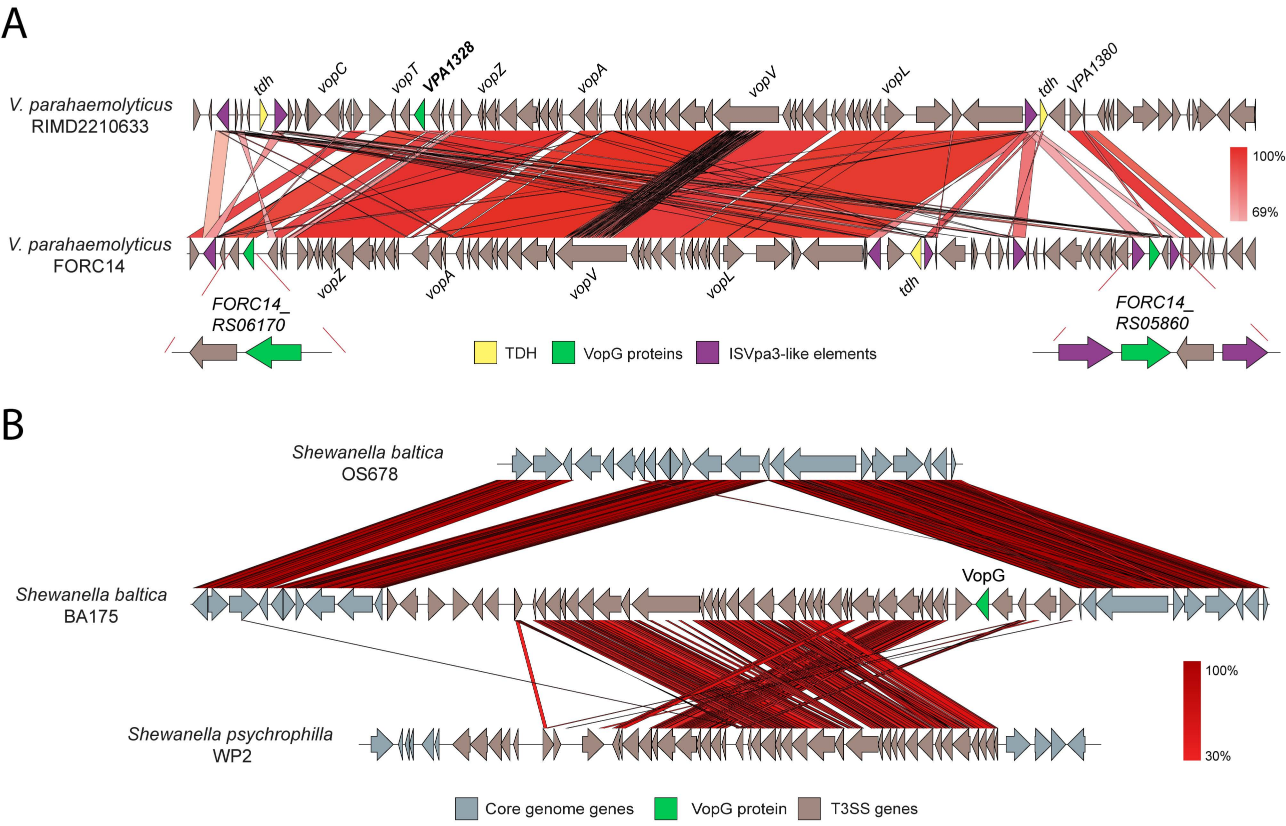

### Figure S3

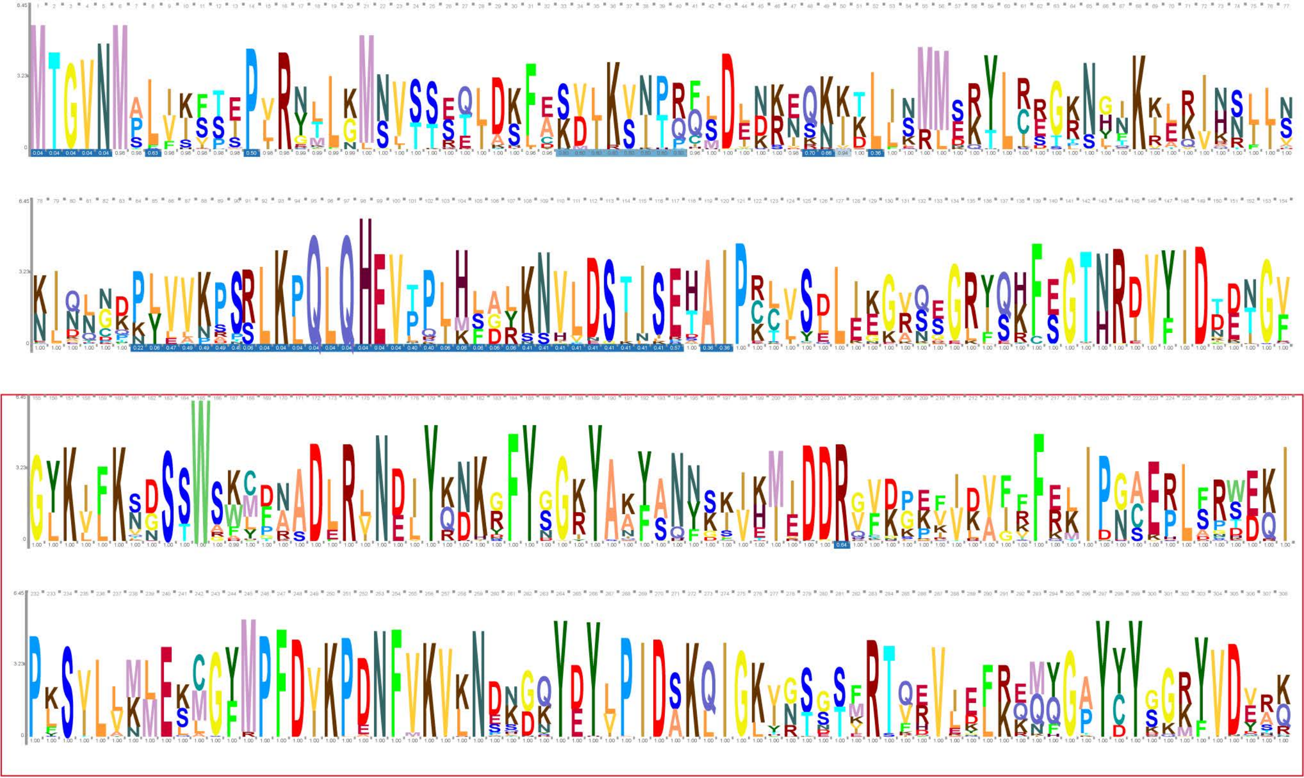

C-terminal region

### Figure S4

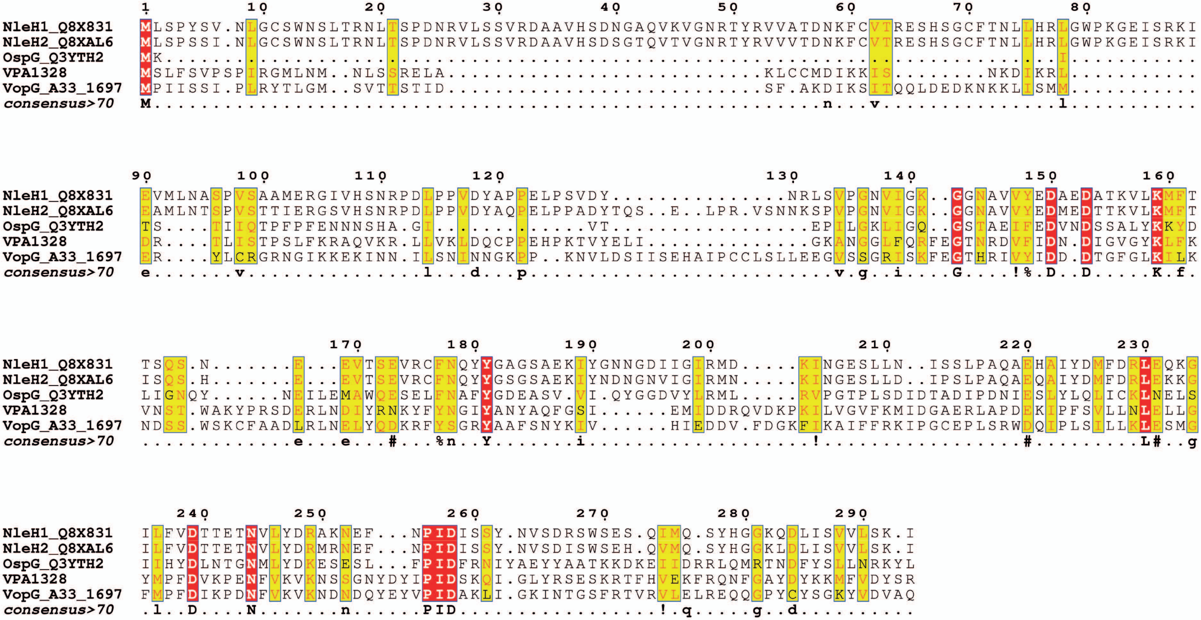

### Figure S5

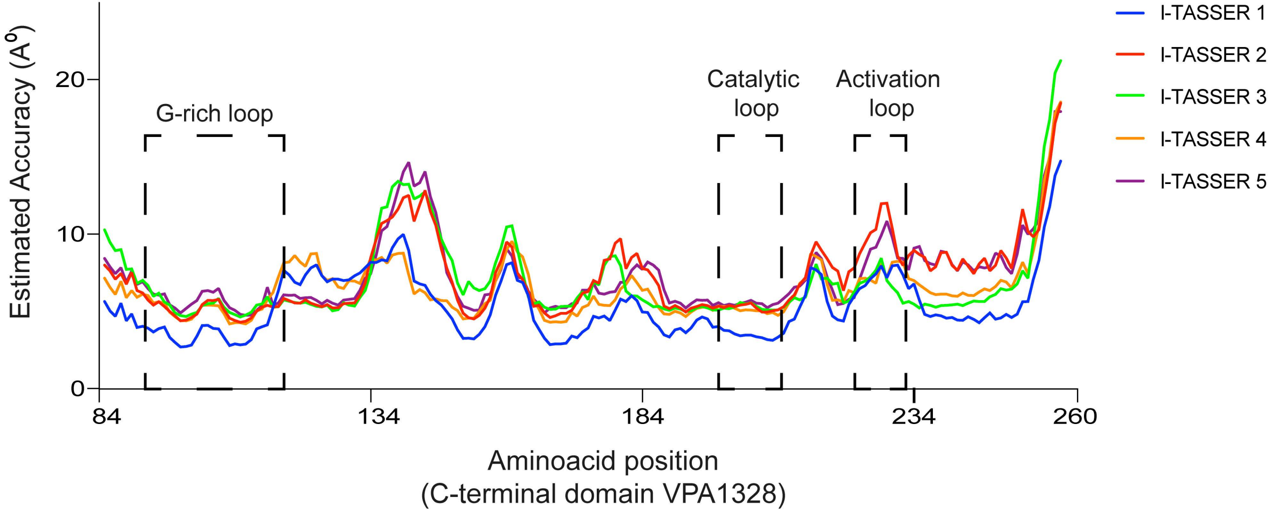
