## Supplementary material for "Identification of a family of *Vibrio* type III secretion system effectors that contain a conserved serine/threonine kinase domain": Figure S2

```
                 1        10        20        30        40        50        60        70        80        90
FORC14_RS06170   ATGTCTTTGTTTTCTGTACCATCTCCTATTAGAGGGATGGTAAAAATGAATTTATCATCTCAAGAACTAGCTAAGCTTTGTTGTATGGAT
FORC14_RS05860   ATGTCTTTAATCCATATTTCCTCTCCTATTCGTAGAATGTTAAACATGAATATCTCATCTCGAGAGTTAGCTGATATCTATCACATGGAT
consensus>70     ATGTCTTT..T...T.T..C.TCTCCTATT.G..G.ATG.TAAA.ATGAAT.T.TCATCTC.AGA..TAGCT.A..T.T.T...ATGGAT

                          100       110       120       130       140       150       160       170       180
FORC14_RS06170   ATAAAGAAAATCTCCAATAAAGATATTAAACGTCTAGATAGGACTTTAATTTCGACTACAAACTTGTTTAAGAGAGCGCAAGTTAAGCGA
FORC14_RS05860   ATTAAAAGCATATCCAATAAAGATATTGAACGTTTAAATAAAACCTTGAAATCGACAAAAAACTTATTTAAGAGAACGAAAGTACAACGA
consensus>70     AT.AA.A..AT.TCCAATAAAGATATT.AACGT.TA.ATA..AC.TT.A..TCGAC.A.AAACTT.TTTAAGAGA.CG.AAGT..A.CGA

                          190       200       210       220       230       240       250       260       270
FORC14_RS06170   CTTTTGGTTAAACTTGACCAATGTCCACCCGAGCATCCTAAAACGGTTTATGAGTTAATTGGTAAAGCCAATGGAGGGTTGTTCCAGCGT
FORC14_RS05860   CTTCTAACTAAAATTGGTTACTGTCAACCAGAACACCCCAAAAGTGTTTGCGAGCTAATTAATAAAGCAAACAGAGGATTACTTCAGCGT
consensus>70     CTT.T...TAAA.TTG...A.TGTC.ACC.GA.CA.CC.AAAA..GTTT..GAG.TAATT..TAAAGC.AA..GAGG.TT..T.CAGCGT

                          280       290       300       310       320       330       340       350       360
FORC14_RS06170   TTTGAAGGAACTAACAGAGACGTATTTATTGATAACGATATTGGTGTTGGATATAAACTATTTAAAGTAAATAGTACATGGGCTAAGTTT
FORC14_RS05860   TTCGAAGGGACGAATCGAGATGTCTTTATTGATAATGAAATTGGATTTGGTTATAAACTATTTAAATTGGATAGCACATGGTCAGATCAC
consensus>70     TT.GAAGG.AC.AA..GAGA.GT.TTTATTGATAA.GA.ATTGG..TTGG.TATAAACTATTTAAA.T..ATAG.ACATGG.C..A....

                          370       380       390       400       410       420       430       440       450
FORC14_RS06170   CCAAGATCCGATGAGCGTCTTAATGATATTTATAGGAATAAATATTTTTATAATGGTAAATATGCTAATTATGCTCAGTTTGATTCTATA
FORC14_RS05860   CCCAAGGCCGATGAAAGATTGAACGATATTTATAAAAACAAAAGTTTTTATAATGGCTATTATGCTCAATATGCTCATTTTGGTTTTATA
consensus>70     CC.A...CCGATGA..G..T.AA.GATATTTATA..AA.AAA..TTTTTATAATGG..A.TATGCT.A.TATGCTCA.TTTG.TT.TATA

                          460       470       480       490       500       510       520       530       540
FORC14_RS06170   GAGATGATTGACGATCGTCAAGTTGACAAGCCTAAAATATTAGTCGGCGTTTTTAAGATGATCGACGGAGCTGAGCGTTTGGCGCCATAC
FORC14_RS05860   GAAATGGTCGATGATCGTCATACTAATAAACCTAAAGTATTAGTTGGAATATTTAGAACAATACCAGGTGCTGAACGTTTAGATCCTAAA
consensus>70     GA.ATG.T.GA.GATCGTCA...T.A.AA.CCTAAA.TATTAGT.GG..T.TTTA..A..AT....GG.GCTGA.CGTTT.G..CC..A.

                          550       560       570       580       590       600       610       620       630
FORC14_RS06170   GAAAAAATCCCATTCTCAGTTTTACTAAATTTAGAATTGTTAGGTTATATGCCTTTTGATGTGAAGCCTGATAATTTTGTCAAAGTAAAA
FORC14_RS05860   GAAAAAATACCATATTCTATTTTAGTAAAGTTGGAACTATTAGGTTATATGCCGTTTGACATTAAGCCTGACAATTTTGTCAAGGTAAAA
consensus>70     GAAAAAAT.CCAT..TC..TTTTA.TAAA.TT.GAA.T.TTAGGTTATATGCC.TTTGA..T.AAGCCTGA.AATTTTGTCAA.GTAAAA

                          640       650       660       670       680       690       700       710       720
FORC14_RS06170   AATTCTGCTGGAAATTATGACTACCTTCCTATTGACTCTAAGCAAATTGGTTTGTACAAAAGTGACTCTAAGAGAACATTTCATGTAGAT
FORC14_RS05860   AACCCTTTGGGTTATTATGATTACCTTCCTATTGATTCTAAACAAATAGGTTTGCATCGAAGTGATTCAGAAAGAACATTTCAAGTAATA
consensus>70     AA..CT...GG..ATTATGA.TACCTTCCTATTGA.TCTAA.CAAAT.GGTTTG.A...AAGTGA.TC..A.AGAACATTTCA.GTA...

                          730       740       750       760       770       780
FORC14_RS06170   AAATTTAGACGAAACTTCGGGGCTTATGATTATAAAAAGATGTTTGTCGATTATTCGCGTTAG
FORC14_RS05860   AATTTTAGAAAAAAATACGGAACTTATTATTATAATAAGAGATTTGTTGACTATCATTGTTAG
consensus>70     AA.TTTAGA..AAA.T.CGG..CTTAT.ATTATAA.AAGA..TTTGT.GA.TAT....GTTAG
```
